# CoExpPhylo – A Novel Pipeline for Biosynthesis Gene Discovery

**DOI:** 10.1101/2025.04.03.647051

**Authors:** Nele Grünig, Boas Pucker

## Abstract

**Background:** The rapid advancement of sequencing technologies has drastically increased the availability of plant genomic and transcriptomic data, shifting the challenge from data generation to functional interpretation. Identifying genes involved in specialized metabolism remains difficult. While coexpression analysis is a widely used approach to identify genes acting in the same pathway or process, it has limitations, particularly in distinguishing genes coexpressed due to shared regulatory triggers from those directly involved in the same pathway. To enhance functional predictions, integrating phylogenetic analysis provides an additional layer of confidence by considering evolutionary conservation. Here, we introduce CoExpPhylo, a computational pipeline that systematically combines coexpression analysis and phylogenetics to identify candidate genes involved in specialized biosynthetic pathways across multiple species based on one to multiple bait gene candidates.

**Results:** CoExpPhylo systematically integrates coexpression information and phylogenetic signals to identify candidate genes involved in specialized biosynthetic pathways. The pipeline consists of multiple computational steps: (1) species-specific coexpression analysis, (2) local sequence alignment to identify orthologs, (3) clustering of candidate genes into Orthologous Coexpressed Groups (OCGs), (4) functional annotation, (5) global sequence alignment, (6) phylogenetic tree generation, and optionally (7) visualization. The workflow is highly customizable, allowing users to adjust correlation thresholds, filtering parameters, and annotation sources. Benchmarking CoExpPhylo on multiple pathways, including the biosynthesis of anthocyanins, proanthocyanidins, and flavonols, as well as lutein and zeaxanthin, confirmed its ability to recover known genes while also suggesting novel candidates.

**Conclusion:** CoExpPhylo provides a systematic framework for identifying candidate genes involved in the specialized metabolism. By integrating coexpression data with phylogenetic clustering, it facilitates the discovery of both conserved and lineage-specific genes. The resulting OCGs offer a strong foundation for further experimental validation, bridging the gap between computational predictions and functional characterization. Future improvements, such as incorporating multi-species reference databases and refining clustering for large gene families, could further enhance its resolution. Overall, CoExpPhylo represents a valuable tool for accelerating pathway elucidation and advancing our understanding of specialized metabolism in plants.

## Background

The continuous advancements in sequencing technologies have led to a drastic reduction in the cost of genome sequencing and the availability of thousands of plant genome sequences [1]. As a result, the primary challenge in genomics has shifted from generating sequencing data to efficiently analyzing and interpreting the vast amount of publicly available data [2]. Databases such as the Sequence Read Archive (SRA) and the National Center for Biotechnology Information (NCBI) provide an extensive collection of sequencing datasets, offering unprecedented opportunities for comparative and functional genomics [3]. While identifying genes structurally becomes more feasible through the integration of transcriptomic data and the facilitated transfer of gene annotations between species through programs such as GeMoMa [4], functionally annotating genes remains a major challenge [5]. In plants, understanding the roles of genes in the complex specialized metabolism is a particular challenge and heavily dependent on information about the biochemical functions of gene products.

One challenge in finding genes belonging to a certain metabolic pathway in plants lies in their scatteredness across the genome. Genes involved in pathways that lead to specialized metabolites are rarely physically clustered [6]. Instead, transcriptional control by a shared transcription factor or transcription factor complex activates all genes involved in a given pathway [7–10]. Notable exceptions are biosynthesis gene clusters, including those reported for withanolide biosynthesis in *Solanaceae*, noscapine biosynthesis in *Papaver somniferum,* or terpenoid biosynthesis pathways such as thalianol, marneral, tirucalladienol, and arabidol in *Arabidopsis thaliana*, among a limited number of other known cases [11–14]. Hence, finding all genes participating in a given pathway based on already known genes merely based on their genomic position can be challenging. Here, coexpression analyses can come to the rescue: Naturally, genes that are required for the same pathway show a similar activity as the encoded enzymes are required simultaneously. This is realized by transcription factors that often regulate the expression of several genes participating in a pathway. Genes whose expression patterns have a correlation to the expression of already known genes of the pathway of interest, might also participate in that pathway due to guilt-by-association [15]. By using Spearman’s rank correlation method, which also detects non-linear correlations, even transcription factors involved in the regulation of a target pathway can be identified. However, the results should be taken with caution, as guilt-by-association has its limits and especially genes that are expressed constantly, such as genes involved in the central metabolism or genes coexpressed due to shared regulatory triggers rather than direct functional relationships can produce misleading results [16]. Over the years, a range of computational strategies has been developed to overcome these challenges. Early network-based approaches such as AraNet v2 leverage cofunctional gene networks to predict pathway membership, but often lack resolution among closely related paralogs or across diverse species, as it contains information only about 28 species [17]. More recently, supervised machine-learning models have been applied to plant pathway gene discovery – for example, a support-vector-machine model trained on over 10 000 genomic and network-based features (conservation, protein domains, duplication events, epigenetic marks, expression and network metrics) to distinguish specialized-from general-metabolism genes in *Arabidopsis thaliana* [18], and RafSee, a Random Forest classifier that integrates sequence-derived, evolutionary, and epigenetic features with coexpression metrics to prioritize candidate genes for specific pathways [19]. These approaches, however, depend on large, carefully curated training sets and do not explicitly incorporate evolutionary context, limiting their applicability to new species or poorly annotated pathways.

To address this limitation, integrating phylogenetic analysis with coexpression data provides a more reliable strategy for identifying functionally relevant genes belonging to one biosynthetic pathway. Phylogenetic approaches help discern evolutionary conservation and functional divergence, enhancing the accuracy of gene function predictions.

A well studied model system is required as proof-of-concept for a new computational approach for the identification of genes in a biosynthesis pathway based on integrated coexpression and phylogenetic integration. The flavonoid biosynthesis with a set of different compounds produced by individual branches is a prime example [20, 21]. Anthocyanins confer colors to fruits and flowers [22], proanthocyanidins (PAs) are responsible for seed coat pigmentation [23, 24] and flavonols are known as UV-response compounds [25, 26] and have been associated with regulatory functions [27]. Due to a plethora of modification reactions catalyzed by numerous (promiscuous) enzymes, there are thousands of different flavonoid derivatives in plants [28, 29]. Despite decades of intense research on the flavonoid biosynthesis [20, 21, 30], there is still potential for new discoveries as demonstrated by recent publications [31, 32].

A second benchmark pathway is the carotenoid biosynthesis, which represents another well-studied yet complex system of plant specialized metabolism. Carotenoids are C40 isoprenoid pigments with essential roles in photosynthesis and photoprotection, as well as in the coloration of flowers and fruits [33]. Lutein and zeaxanthin, two major xanthophylls, are prominent products of this pathway. The extensive research and the identification of many core enzymes [34–36] makes carotenoid biosynthesis, alongside flavonoids, an excellent model for benchmarking computational approaches for gene identification.

Here, we present CoExpPhylo as a novel implementation for automatic integration of coexpression and phylogenetic analyses for the identification of players in a biosynthesis pathway. Starting from minimal prior knowledge about some genes involved in the biosynthesis of a metabolite of interest, CoExpPhylo enables the identification of candidate genes associated with the biosynthesis of that metabolite. The potential is demonstrated by analyses on the biosynthesis of the flavonoids anthocyanin, proanthocyanidin, and flavonol as well as the carotenoids lutein and zeaxanthin.

## Implementation

CoExpPhylo was written in Python3 using SciPy v1.13.0 [37], NumPy v1.26.4 [38], Networkx v3.3 [39] and Plotly v5.22.0 [40] as well as the utility GNU parallel 20210822 [41] through bash scripting. CoExpPhylo utilizes different external tools: Several local alignments are conducted via DIAMOND v2.0.14.152 [42], global alignments are executed using MAFFT v7.490 [43, 44] or optionally by MUSCLE v3.8.1551 [45] and phylogenetic trees are inferred by FastTree v2.1.11 [46, 47], RAxML-NG v1.2.2 [48] or IQ-TREE v2.0.7 [49]. Optionally, the final tree files can be uploaded to iTOL [50].

### Input data collection

The analysis is conducted based on transcriptomic datasets of 240 species from various orders (Figure 1, Additional File 1) [51, 52]. Briefly, the coding sequences were collected from Phytozome (https://phytozome-next.jgi.doe.gov), NCBI GenBank (https://www.ncbi.nlm.nih.gov/genbank), and species-specific websites. The RNA-seq data was collected from the NCBI SRA (https://www.ncbi.nlm.nih.gov/sra) [3] using fastq-dump (https://github.com/ncbi/sra-tools) and processed as described in the methodology of the dataset publications: Count tables and gzip-compressed FASTQ files were processed using kallisto v0.44 [53] and subsequently merged into a single count table per species with custom Python scripts [54]. All values in the count tables are Transcripts Per Million (TPM). Samples were excluded, if the total number of reads was below one million or the read distribution did not match RNA-seq expectation i.e. at least 20% of reads assigned to the most abundant 100 transcripts. This was handled using the Python script filter_RNAseq_samples.py v0.4 (https://github.com/bpucker/CoExp). Each annotation file was generated with the Python script construct_anno.py v0.1 (https://github.com/bpucker/PBBtools). For each dataset, the gene IDs in each file were cleaned as any special characters were replaced. Only one representative transcript was retained per gene by selecting the one with the longest coding sequence (CDS). This established approach was chosen to reduce redundancy in the dataset to limit computational complexity. The isoform with the longest CDS often corresponds to the most functionally relevant isoform [55]. In cases where transcript identifiers were missing from the sequence headers (e.g. no transcript number or suffix indicating alternative isoforms), transcripts belonging to the same gene were determined and reduced via isoform_purger.py v0.21 (https://github.com/bpucker/PBBtools).

**Figure 1:**
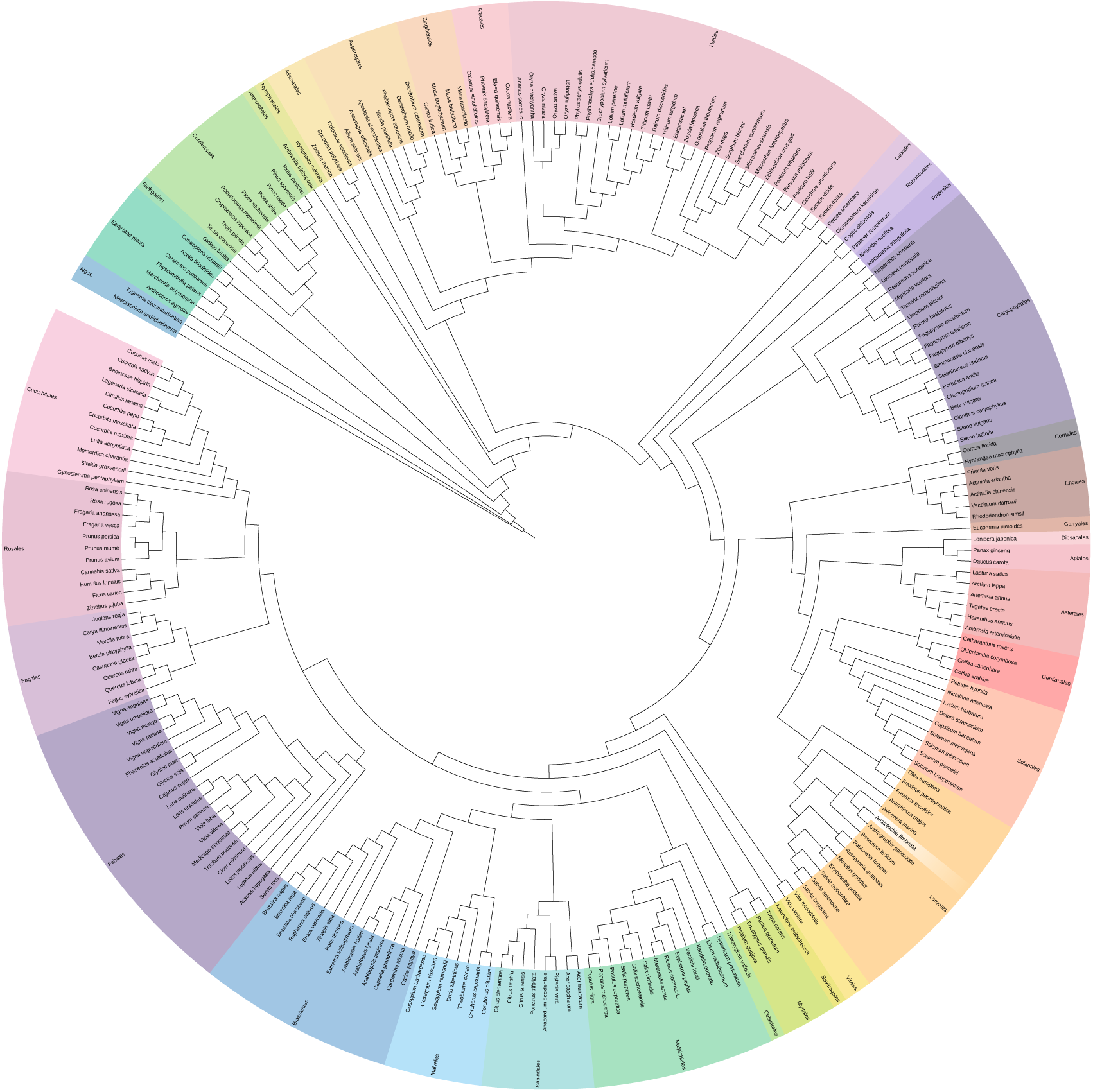
Taxonomic grouping of species included in this study. Colored ranges represent the taxonomic orders of the species. For better readability, the classification of early land plants and *Coniferopsida* has been simplified. *Aristolochia fimbriata* appears to be misplaced, as it belongs to the *Piperales*, which would be placed close to the *Laurales*. The figure was generated using iTOL [50].

The CoExpPhylo workflow is designed to start from a known metabolite or biosynthetic pathway and a small set of bait genes that are already associated with this pathway. Using these starting points, CoExpPhylo identifies additional candidate genes that might be involved in the same biosynthesis process. To provide the necessary information for this analysis, a configuration (config) file is required. This file defines all datasets (i.e., species) to be included and specifies the paths to the relevant input data. The columns in this file need to be comma-separated without a header. Each row describes one dataset (i.e. one species). The columns contain (species-)IDs, path to the species-specific count table, path to a species-specific multiple FASTA file containing coding sequences, and the path to the species-specific bait file (Table 1). An example config file as well as a synthetic sample dataset demonstrating the required input formats are available in the GitHub repository (https://github.com/bpucker/CoExpPhylo).

**Table 1:** Explanation and examples of the columns in the config file needed for CoExpPhylo.

| ID | Path to TPM file | Path to CDS file | Path to bait file |
| --- | --- | --- | --- |
| Unique; usually the species name, is later used as suffix | Path to a count table, first column of count table includes gene IDs, first row contains sample names, other fields are expression values | Path to a multiple FASTA file with coding sequences. Sequence IDs must match sequence IDs in count table | Path to text file with initial bait sequence IDs. IDs must match sequence IDs in TPM and CDS file; one sequence per line |
| *Arabidopsis-thaliana* | /path/to/Athaliana.tpm.tsv | /path/to/Athaliana.cds.fasta | /path/to/Athaliana.baits_anthos.txt |

A key element of the CoExpPhylo workflow are the baits, which define the set of genes already known to be involved in the biosynthetic pathway of interest. Depending on the available knowledge, this set can consist of multiple genes or even a single gene that serves as the starting point for the analysis. If possible, we recommend selecting late players of the pathway – genes that act close to the final product. Ideally, these bait genes should be specific to the pathway rather than shared with other metabolic branches, and orthologous genes should be provided for each species included in the study. The bait file must be a simple text file, with one sequence ID per line.

To annotate each OCG properly, Araport11 *Arabidopsis thaliana* Col-0 reference polypeptide sequences and annotation terms derived from TAIR10 and Araport11 [56, 57] were used.

## Results

The CoExpPhylo workflow systematically analyzes coexpressed genes within one biosynthetic pathway across multiple species, integrating coexpression results, sequence alignment, clustering, and phylogenetic relationships. The following steps outline the approach, with a schematic representation provided in Figure 2.

**Figure 2:**
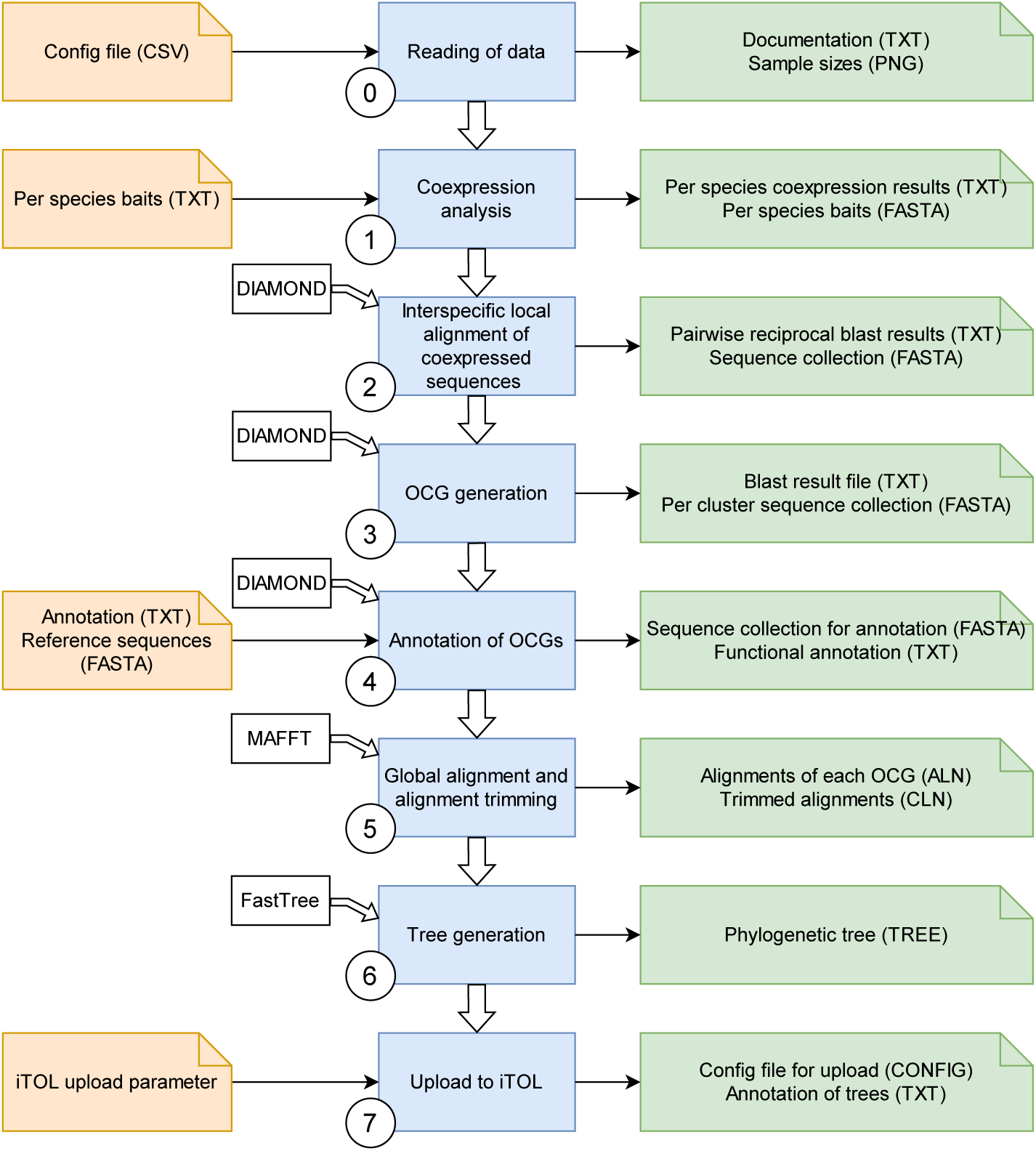
Workflow of CoExpPhylo. Shown is a schematic overview of the pipeline of coexp_phylo.py. The workflow steps are represented in blue, input files and their file extension are denoted in orange, whereas the output files and their file extensions are colored in green. White boxes indicate the utilization of external tools. OCG means Orthologous Coexpressed Group. This flow chart was generated with drawio.com.

### Step 1 – Coexpression analysis

After loading of the input data (Step 0), the coexpression is conducted for every species separately. The IDs of the genes for which the coexpression analysis shall be conducted are loaded from the respective baits file. To evaluate the correlation between the gene expression of the baits and the remaining genes, Spearman’s rank correlation was applied *via* the spearmanr-function from the module SciPy [37] in Python to calculate the Spearman’s rank correlation coefficient *r*s and the corresponding adjusted p-value. The default correlation coefficient cut-off is set to 0.7 but can be adjusted using the argument `--r`. The p-value threshold can be adjusted by specifying the `–p` argument. If not set, the widely accepted default value of 0.05 is used. To further customize the output of the coexpression analyses according to the structure of the input dataset, two additional parameters can be adjusted: Through specification of the argument `--numcut`, the maximal number of genes that are retrieved from each co-expression analysis can be lowered or increased. The default value is set to 100, therefore, only the top 100 sequences found *via* coexpression analysis are considered for computation in further steps. Lowering the value can reduce the number of relevant genes fetched from the analyses, whereas increasing the value will substantially extend the run time. Secondly, `- -min_exp_cutoff` specifies the minimal cumulated expression of any gene that is detected *via* coexpression analysis among all samples. The default is set to 30, which can help to exclude technical artifacts.

### Step 2 – Local alignment of coexpressed sequences

Candidate sequences are identified *via* DIAMOND [42]. All sequences collected in the coexpression analyses employed as query, with every species serving as a database. This requires several local alignment runs to be conducted. DIAMOND blastp runs with default parameters, the results are stored in a tabular output format 6. Promising candidate sequences can then be filtered *via* different parameters and are written into one FASTA file. Candidate sequences must exceed the following requirements: Firstly, the e-value of the alignment must not exceed 10-5. Secondly, the bit score must be greater than 100, thirdly, the alignment length must surpass 100, and fourthly, the similarity must be greater than 80%. Those values can be adjusted by setting `--evalue`, `--scorecut`, `--lencut` as well as `--simcut`, respectively. The influence of lowering the length or similarity score cutoff is shown in Additional File 2.

### Step 3 – Generation of Orthologous Coexpressed Groups (OCGs)

To divide the sequence collection into clusters, another DIAMOND blastp run is required. At this time, the sequence collection was employed as query and simultaneously as database. The results were filtered according to the previously described parameters. The remaining pairs are incorporated into graphs using the Graph function from the Python module NetworkX [39]. Every sequence that occurs in the filtered BLAST result file is added to the graph as a node, whereas edges are drawn between sequences that have a high blastp result and thus, appear in the filtered blast-result file. Thereby, all sequences with a high sequence similarity are connected within one graph. Finally, all sequences of each OCG are written into one FASTA file per OCG. The OCGs are ranked according to the occurrence of sequences derived from the coexpression analysis. Hence, OCGs with a high ratio between sequences identified *via* coexpression analysis and orthologs identified *via* DIAMOND blastp are labeled firstly, as they presumably contribute more profoundly to the pathway as OCGs with a low number of coexpressed sequences. OCGs with less than ten sequences in general are excluded as well as OCGs that contain coexpressed sequences from less than three species. Thus, the results have a higher significance, as artifacts are excluded.

### Step 4 – Functional annotation of OCGs

Optionally, OCGs can be annotated to gauge functionality of the genes within one OCG. To enable this functional annotation, a reference peptide FASTA file must be provided by using `- -reference`. If the annotation of the OCGs should not only contain the sequence ID but also a functional description, an annotation table can be specified *via* `--anno`. This annotation file must be a tab-separated table with the first column containing matching IDs to the provided FASTA file and the second column comprises the respective function. After the sequences collected in step 4 are clustered, a defined percentage of sequences is collected per OCG and written into a separate FASTA file. The percentage can be defined with `-- seqs_cluster_anno` and is assigned to 50 by default. Thus, 50% of the sequences of each OCG are selected randomly and are chosen for the annotation in order to save computational resources. However, to ensure the assignment of an annotation valid for the majority of sequences in the OCG, the value cannot be set to less than 10%. If the value is lower, it will automatically be defined as 10, representing 10% of the sequences. Additionally, at least five sequences are employed for the annotation. Howsoever, to annotate an OCG properly, the sequences of each OCG are locally aligned *via* DIAMOND [42] against a database built from the reference FASTA file. Subsequently, the best hit of each alignment is retrieved according to the bit score. Among all annotations of each OCG, the most common one is selected as the functional annotation for the respective OCG. To elucidate the accuracy of the annotation, a reliability score is displayed in the functional annotation table. This score indicates the proportion of sequences that were annotated with the conclusively selected annotation.

### Step 5 – Global alignment

The global alignment of each OCG is executed either by MAFFT [43, 44] or optionally with MUSCLE [45]. With the argument `--alnmethod`, the tool can be selected. By default, MAFFT is used. If the executable for the intended tool is not included in the user’s system PATH environment variable, the path to the tool must be provided with `--mafft` or `-- muscle`, respectively. The global alignment is executed using the default parameters of the tools. Afterwards, the aligned sequences are trimmed, so that only positions with sufficient occupancy are kept. The occupancy cutoff is set to 10% but can be adjusted by the user with the argument `--occupancy`. The output file is written in FASTA format.

### Step 6 – Phylogenetic tree generation

The cleaned and trimmed alignment files are subsequently used as input files for the phylogenetic tree construction. With the argument `--treemethod`, the tool for tree generation can be specified. By default, FastTree [46, 47] is utilized with the options `-wag` and `-nosupport`. Alternatively, RAxML-NG [48] using the model “LG+8G+F” or IQTREE [49] can be applied for this step. To enable the upload to common phylogenetic tree visualization programs, especially the batch upload to iTOL [50], the output files have Newick format.

### Step 7 – Batch upload to iTOL

After the CoExpPhylo analysis finished, the tree files can be uploaded to any phylogenetic tree visualization software. As the number of generated tree files can be complex to handle manually, it is possible to upload the tree files automatically to iTOL [50]. This upload requires a Perl script that can be downloaded from the iTOL help page (https://itol.embl.de/help.cgi#batch). By using the argument `--upload_script`, the path to this upload script must be provided if it is not stored in the same folder as the main Python script coexp_phylo.py. Furthermore, an annotation file is uploaded with every tree that enables coloration of the sequences tagged with the suffix “_coexp”, i.e. sequences that were collected *via* coexpression analyses, in iTOL. The automatic batch upload is only possible with an active standard subscription and requires the specification of the API using the argument `--API`. All trees are uploaded into the project that is defined by `--pro_name`. This project must already exist in the user’s account and should be unique among all workspaces. To avoid failure of the script due to misspelling in the API or project name, a test-tree is uploaded during the initial step if a batch upload to iTOL is desired. An error during the test upload leads to the termination of the whole coexp_phylo.py script with a corresponding error message.

### Output files

CoExpPhylo generates several final output files summarizing the analysis results (Table 2). The documentation file records the script version, parameter settings, external tool versions, and input file paths, including an MD5 hash for each file. To assess species representation, an HTML-based species count histogram visualizes the distribution of coexpressed sequences per species. This allows users to detect potential biases, such as underrepresented species with low coexpression signal due to limited sample availability.

**Table 2:**
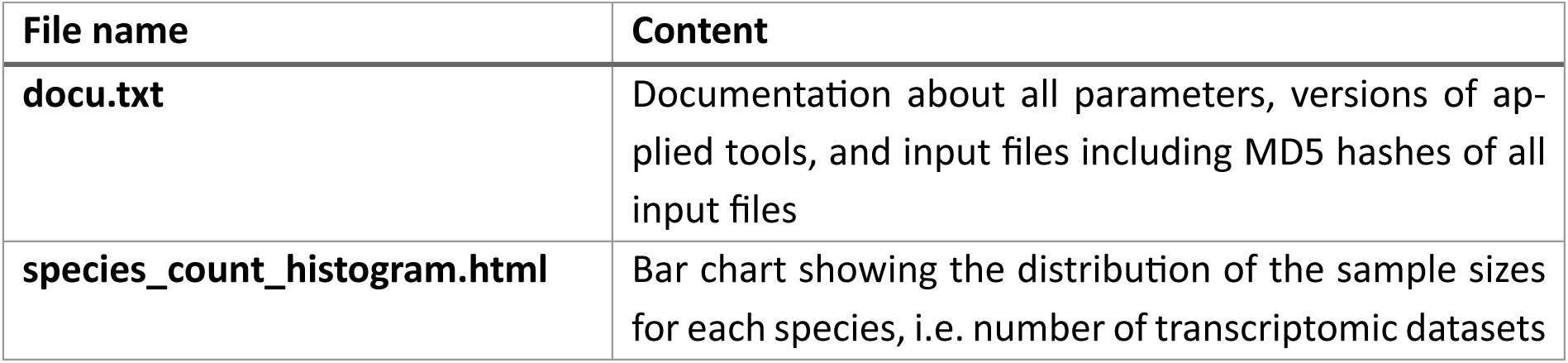
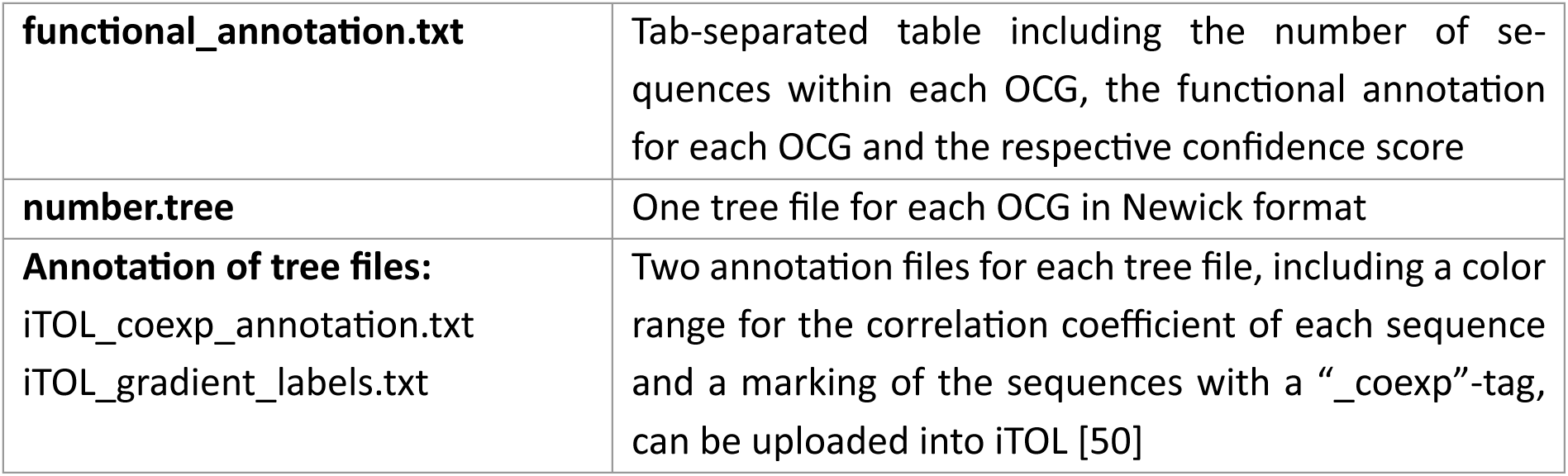
Overview over the core output files generated by CoExpPhylo.

| File name | Content |
| --- | --- |
| **docu.txt** | Documentation about all parameters, versions of applied tools, and input files including MD5 hashes of all input files |
| **species_count_histogram.html** | Bar chart showing the distribution of the sample sizes for each species, i.e. number of transcriptomic datasets |
| **functional_annotation.txt** | Tab-separated table including the number of sequences within each OCG, the functional annotation for each OCG and the respective confidence score |
| **number.tree** | One tree file for each OCG in Newick format |
| **Annotation of tree files:**<br>iTOL_coexp_annotation.txt<br>iTOL_gradient_labels.txt | Two annotation files for each tree file, including a color range for the correlation coefficient of each sequence and a marking of the sequences with a “_coexp”-tag, can be uploaded into iTOL [50] |

A tab-separated ranking table provides an overview of OCGs based on the proportion of co-expressed sequences. This table includes the number of sequences per OCG, annotation details (if applied), and a confidence score reflecting the relationship to the assigned annotation sequence. The confidence score is determined by comparing a subset of sequences from each OCG to a reference database, reflecting the proportion of sequences with high similarity to the final annotation sequence.

For phylogenetic analysis, a Newick-formatted tree file is generated for each OCG, representing the evolutionary relationships among the identified sequences (Figure 3). Additionally, two annotation files are produced per OCG to facilitate visualization in iTOL [50] if automatic upload is not enabled. The first annotation file (*iTOL_coexp_annotation.txt*) highlights sequences identified through the initial intra-specific coexpression analysis. The second annotation file (*iTOL_gradient_labels.txt*) assigns a color bar to each sequence, reflecting its highest correlation coefficient. This visualization aids in assessing the overall correlation strength of an OCG to the biosynthetic pathway under investigation.

**Figure 3:**
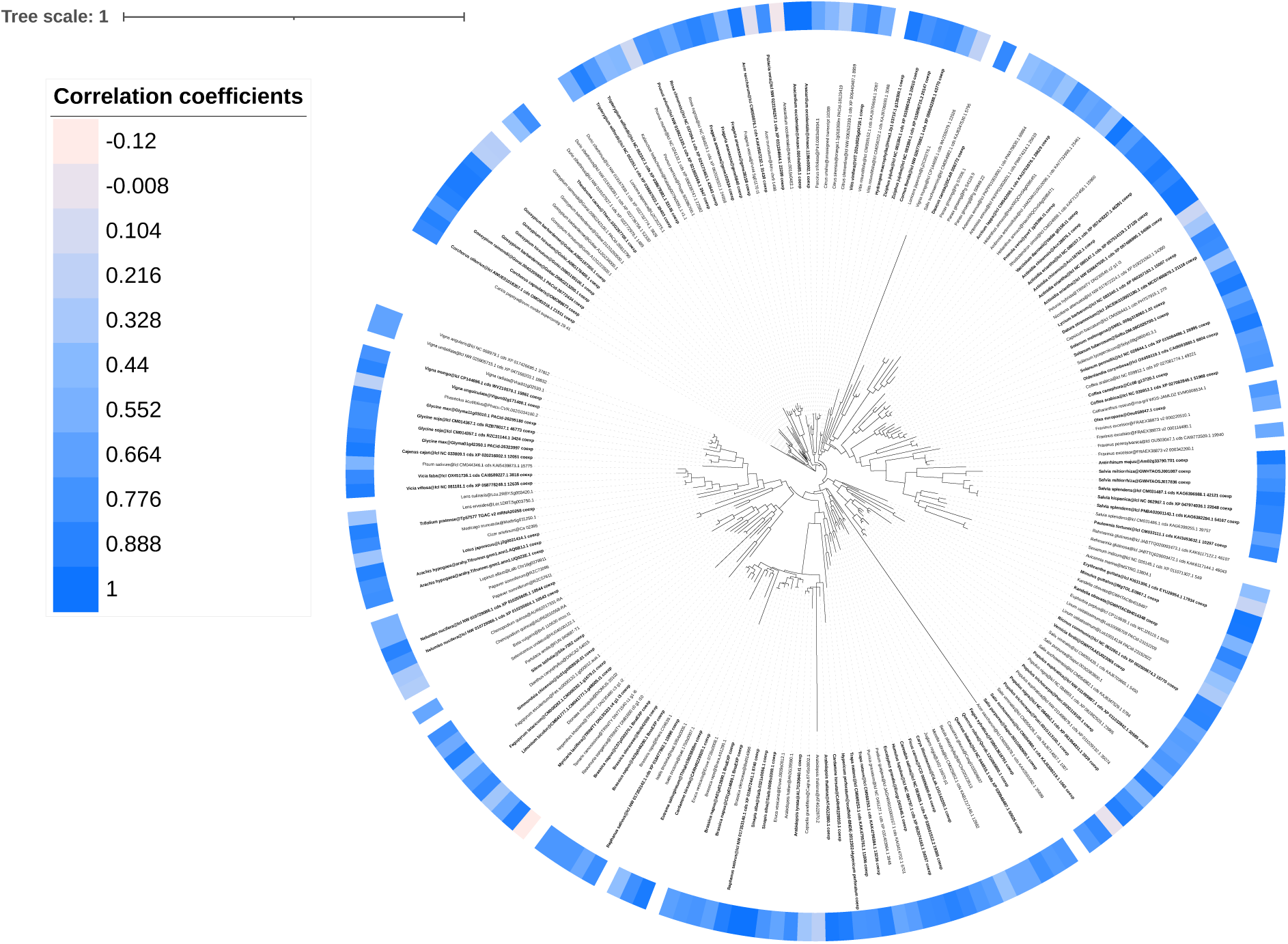
Exemplary representation of the visualization of one Step via iTOL. [**50**]. Shown is the result of Orthologous Coexpressed Group (OCG) 0009 retrieved through CoExpPhylo with respect to the anthocyanin biosynthesis synthesis. CoExpPhylo was conducted based on the transcriptomic data of 239 species [51, 52], dihydroflavonol 4-reductase (DFR), anthocyanidin synthase (ANS), and the anthocyanin-related glutathione S-transferases (arGSTs) Bronze2 (Bz2) as well as transparent testa 19 (TT19) were used as baits. Candidate baits were identified via KIPEs3 [58, 59]. Any species without identifiable baits as well as samples with a high expression (> 5 tpm) of leucoanthocyanidin reductase (LAR) or anthocyanidin reductase (ANR) were excluded from the analysis. The global alignment was conducted with MAFFT [43, 44]. The trees were constructed with FastTree [46, 47].

### Proof of concept: application to flavonoid biosynthesis

To evaluate the performance, we applied CoExpPhylo to three distinct branches within the flavonoid biosynthesis: anthocyanins, PAs, and flavonols. We used respective genes as input baits and were able to detect various OCGs annotated with already known players of the biosynthetic pathway as well as possible candidate genes (Figure 4A).

**Figure 4:**
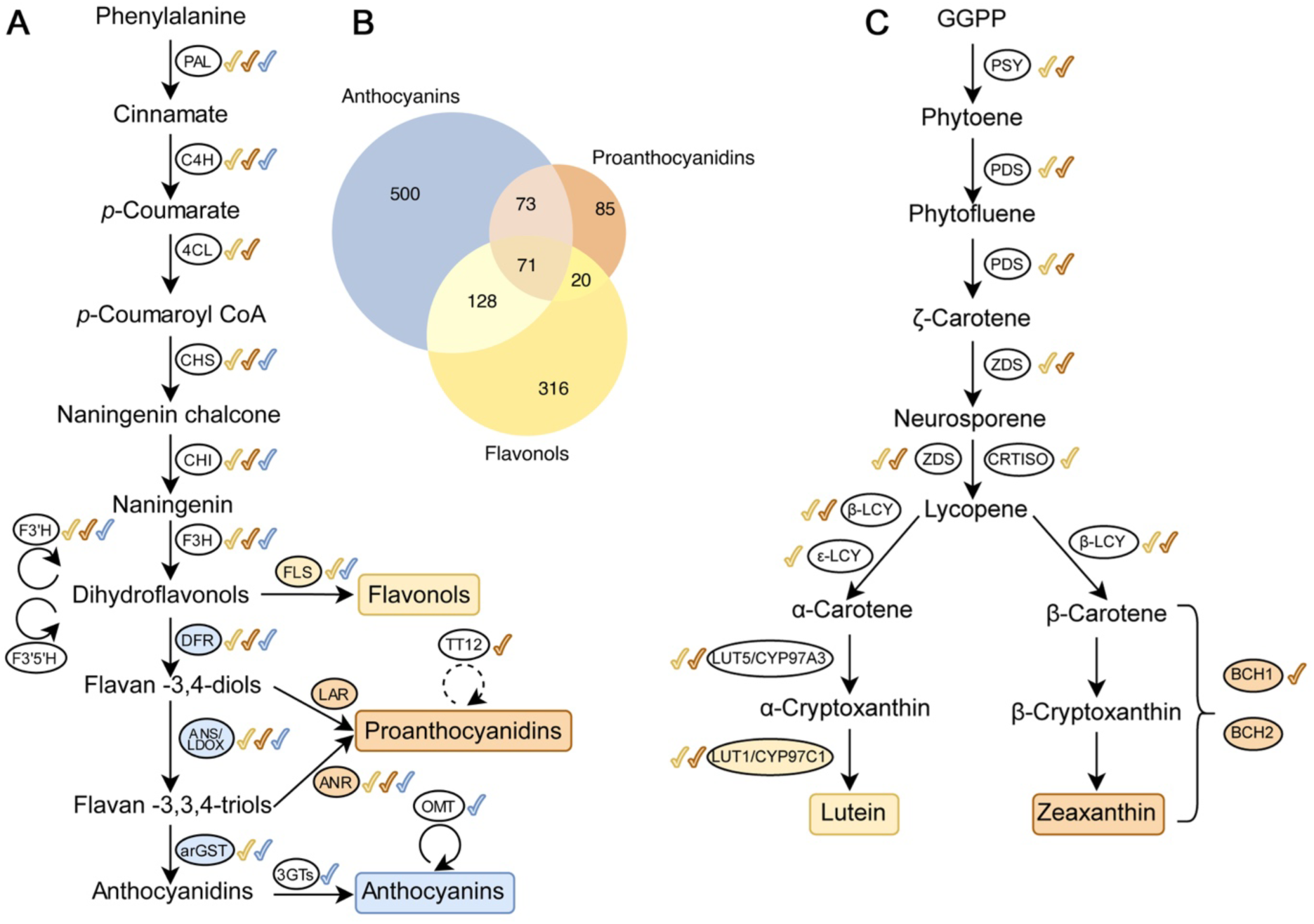
Schematic overview of different branches of the flavonoid and carotenoid biosynthesis and the detection via CoExpPhylo. Enzymes highlighted with a colored background were used as input baits for the analysis of the corresponding biosynthetic branch. Colored check marks indicate the Orthologous Coexpressed Groups (OCGs) retrieved for that branch. **(A)** Displayed enzymes are PHE ammonia lyase (PAL), cinnamate 4-hydroxylase (C4H), 4-coumarate:CoA ligase (4CL), chalcone synthase (CHS), chalcone isomerase (CHI), flavanone 3-hydroxylase (F3H), flavonoid 3’-hydroxylase (F3’H), flavonoid 3’,5’-hydroxylase (F3’5’H), flavonol synthase (FLS), dihydroflavonol 4-reductase (DFR), leucoanthocyanidin reductase (LAR), transparent testa 12 (TT12), anthocyanidin synthase/leucoanthocyanidin dioxygenase (ANS/LDOX), anthocyanidin reductase (ANR), anthocyanin-related glutathione S-transferase (arGST), UDP-dependent anthocyanidin-3-O-glycosyltransferase (3GT), O-methyltransferase (OMT). The dotted arrow indicates a transporter. **(B)** Venn diagram showing the overlap of genes identified in the three biosynthetic pathways: anthocyanins, proanthocyanidins and flavonols. **(C)** Displayed enzymes are phytoene synthase (PSY), phytoene desaturase (PDS), ζ-carotene desaturase (ZDS), carotenoid isomerase (CRTISO), lycopene β -cyclase (β-LCY), lycopene ε -cyclase (ε-LCY), lutein deficient5/cytochrome P450-family 97-A3 (LUT5/CYP97A3), lutein deficient1/cytochrome P450-family 97-C1 (LUT1/CYP97C1), beta carotenoid hydroxylase 1 (BCH1), and beta carotenoid hydroxylase 2 (BCH2).

Briefly, flavonoid biosynthesis originates from the general phenylpropanoid pathway, which involves the key enzymes phenylalanine ammonia-lyase (PAL), cinnamate 4-hydroxylase (C4H), and 4-coumarate:CoA ligase (4CL). The core flavonoid biosynthesis pathway includes chalcone synthase (CHS), chalcone isomerase (CHI), and flavanone 3-hydroxylase (F3H). The emerged dihydroflavonols can be further diversified through the flavonoid 3’-hydroxylase (F3’H) and the flavonoid 3’,5’-hydroxylase (F3’5’H) [20, 60]. Flavonol biosynthesis proceeds via flavonol synthase (FLS), while the anthocyanin biosynthesis requires dihydroflavonol 4-reductase (DFR), anthocyanidin synthase/leucoanthocyanidin dioxygenase (ANS/LDOX), anthocyanin-related glutathione S-transferase (arGST), and UDP-dependent anthocyanidin-3-O-glycosyltransferase (3GT) [20, 31]. PAs can be synthesized through leucoanthocyanidin reductase (LAR) or anthocyanidin reductase (ANR).

The flavonoid biosynthesis is transcriptionally regulated by a complex consisting of a MYB, a bHLH, and a WD40 transcription factor, collectively forming an MBW complex [9, 61]. The MYB component determines the target gene, forming a complex with bHLH42 and TTG1 [62]. MYB11, MYB12 and MYB111 regulate the flavonol biosynthesis bHLH-independent [7], while proanthocyanidin biosynthesis is regulated through MYB123 in a complex with a bHLH and a WD40 component [63]. MYB75, MYB90, MYB113, and MYB114 are responsible for anthocyanin biosynthesis regulation [9, 64, 65].

#### Anthocyanins

Focusing on the anthocyanin branch, which is responsible for the vibrant pigmentation in flowers and fruits, late-stage enzymes were selected as bait candidates to evaluate CoExpPhylo’s capability in identifying genes involved in specialized metabolic pathways. This approach aimed to determine whether the tool could accurately recover established biosynthetic genes and differentiate them from those associated with related flavonoid branches.

The final enzymatic step in anthocyanin biosynthesis is catalyzed by UDP-glucose:flavonoid glucosyltransferases (UFGTs). However, UFGTs cannot yet be reliably annotated due to the limited knowledge about their conserved amino acid residues. To avoid introducing uncertainty into the analysis, UFGTs were excluded from the bait selection. As the lack of a comprehensive set of amino acid residues required for the activity of arGSTs complicates their identification across species, DFR and ANS/LDOX were also included as baits to ensure a robust test case for CoExpPhylo.

A major challenge in using *DFR* and *ANS* as baits is their dual involvement in anthocyanin and PA biosynthesis. To mitigate potential misassignments, RNA-seq samples with considerable expression (>5 tpm) of either LAR or ANR – late-stage enzymes of PA biosynthesis – were excluded. Since low expression of LAR and ANR suggests minimal PA biosynthesis, the remaining DFR and ANS expression data are more likely linked to anthocyanin biosynthesis. By applying these selection criteria, we aimed to create a controlled test scenario that allows for an initial evaluation of CoExpPhylo’s ability to distinguish biosynthetic genes from related pathways and to identify potential new candidates.

The output (Additional File 3) shows OCGs associated with multiple key enzymes of the anthocyanin biosynthesis. One OCG was annotated with PAL and another with C4H, both enzymes of the phenylpropanoid pathway. Additionally, CHS, CHI, F3H and F3’H were assigned to distinct sequence clusters. Interestingly, no OCG corresponding to 4CL, which acts between C4H and CHS in the pathway, was detected. This absence could be due to the gene’s diverse functional roles beyond flavonoid biosynthesis its broad substrate specificity (e.g., 4-coumaric acid, ferulic acid, and caffeic acid), or potential limitations in coexpression-based detection [66, 67]. The involvement of 4CL in multiple metabolic branches may result in a distinct expression pattern that does not correlate strongly with flavonoid-specific genes. Due to the phylogenetic divergence of the input species, multiple OCGs were generated for certain genes. For instance, F3H was represented by three OCGs (0017, 0037, and 0069), corresponding to different taxonomic groups: one cluster contained sequences from *Eudicotyledons*, another from *Coniferopsida* and the third from species belonging to the order *Poales*.

The initial bait genes (*DFR*, *ANS*/*LDOX,* and *arGSTs*) were also retrieved as OCGs. Furthermore, the OCGs annotated as UDP-dependent glycosyltransferase UGT78D2, along with the anthocyanin-decorating enzymes SCPL10, AT1, and O-methyltransferase 1 (OMT), were identified. Two additional OCGs (0072 and 0145) were annotated with UDP-dependent glycosyltransferases UGT80B1 and UGT84A1, respectively. Although these enzymes have not been broadly characterized in anthocyanin modification, the *Medicago truncatula* homolog *Mt*UGT84A1 has been reported to be linked to increased anthocyanin accumulation alongside its role in shoot growth [68]. Further analysis of the sequences within these OCGs may reveal additional glycosyltransferases pertinent to anthocyanin biosynthesis. However, the annotation process is based on *Arabidopsis thaliana* reference sequences, which may explain certain limitations. Genes or functions absent in *A. thaliana* cannot be annotated correctly, so that the annotation of UGT80B1 and UGT84A1 might be an artifact thus suggesting that some other UGT lineages could contribute to the diversity of anthocyanins. Similarly, a potential OCG containing sequences encoding F3’5’H was not identified as *A. thaliana* lacks this enzyme, highlighting the importance of a multi-species reference database for the annotation.

Additionally, transcription factors associated with anthocyanin biosynthesis regulation, including MYB90, MYB12, and the bHLH transcription factor bHLH42/TT8, were identified. Notably, FLS was also detected, despite its primary role in flavonol biosynthesis rather than anthocyanin production. However, FLS; F3H and ANS/LDOX belong to the 2-oxoglutarate-dependent dioxygenase (2ODD) superfamily whereby functional redundancy might occur [69].

#### Proanthocyanidins

PAs, also known as condensed tannins, contribute to plant defense and seed coat pigmentation. Their biosynthesis shares multiple enzymatic steps with anthocyanin biosynthesis, making it challenging to delineate genes specific to this pathway. To assess CoExpPhylo’s ability to resolve this complexity, late stage PA biosynthetic genes, including *LAR* and *ANR* (also known as *BANYLUS* (*BAN*)) were used as baits.

The initial bait gene *BAN* was detected three times among the final OCGs. As *A. thaliana* does not have a LAR, no OCG was annotated accordingly. However, all genes encoding enzymes involved in the straight PA biosynthesis, starting with PAL, were successfully retrieved within annotated OCGs (Figure 4A). Notably, genes associated with closely related pathways, such as arGSTs and FLS), were not retrieved, suggesting a certain specificity of CoExpPhylo in distinguishing metabolic branches. Nonetheless, there is a moderate overlap between the analyzed pathways (Figure 4B). This can be explained as the metabolites share the majority of their upstream pathway and by the promiscuity of several enzymes, such as FLS, ANS, and multiple UFGTs [70–74]. Hence, the identification of the same candidate genes in multiple pathways does not indicate any errors, but is biologically plausible and expected.

Additionally, one OCG was annotated with the bHLH transcription factor bHLH42/TT8, which is involved in PA biosynthesis. The main MYB transcription factor regulating PA biosynthesis, MYB123, was not identified as a separate OCG. This highlights a limitation of CoExpPhylo in resolving individual lineages of large gene families such as MYB transcription factors. Due to high sequence similarity within these families, the delineation of OCGs may be influenced by the chosen similarity threshold, potentially leading to the merging of closely related sequences. Possibly, the OCG annotated with MYB5 (0058) contained MYB123 sequences along with other members of MYB subgroup 5. To achieve a more precise resolution of large transcription factor families, dedicated tools like MYB_annotator might show superior performance [75]. Furthermore, the WD40 transcription factor TTG1 was not captured as an OCG. This may be due to its diverse functional roles, spanning various developmental processes and specialized metabolism [63, 76–79] which lead to a constant expression pattern that is not suitable for detection via coexpression analysis.

#### Flavonols

Flavonols, a subclass of flavonoids, function as antioxidants and UV protectants in plants. They serve as precursors for multiple downstream metabolic pathways, making their biosynthetic gene network particularly intricate. To investigate the flavonol branch, FLS, a key enzyme converting dihydroflavonols into flavonols, was selected as a bait sequence. Since FLS competes with DFR for its substrate [80–82], its expression pattern provides a means to distinguish flavonol biosynthesis from related pathways.

All core genes of the flavonol biosynthesis pathway were retrieved within annotated OCGs. However, additional OCGs were identified containing sequences annotated with genes typically associated with anthocyanin biosynthesis, such as *DFR* and *ANS*/*LDOX*, or PA biosynthesis (e.g. *BAN*) which do not directly participate in flavonol production. This suggests that the separation of the flavonol and anthocyanin branches is not entirely distinct in the coexpression analysis, potentially reflecting their shared metabolic precursors and regulatory elements. Moreover, this overlap might be influenced by the diversity of the cell types included in the transcriptomic data. Previous studies have demonstrated cell type-specific expression patterns for different branches of flavonoid biosynthesis [83, 84].

In addition to enzymatic genes, transport and modification-related genes were identified. The transporter DTX35, implicated in flavonol transport, was retrieved as an OCG. Moreover, UDP-dependent glycosyltransferases UGT78D2, UGT84A1, and UGT84A2 were identified, suggesting potential roles in flavonol glycosylation. UGT78D2, an anthocyanin-glycosyltransferase, is present in *Caryophyllales* – an order lacking anthocyanins but having flavonols [85]. This underlines their potential involvement in flavonols glycosylation, which was already reported in *Brassica napus* [86]. Regarding transcription factors, the bHLH regulator bHLH42/TT8 and MYB12 were captured as OCGs. However, MYB11 and MYB111, which together with MYB12 constitute subgroup 7 of the R2R3-MYB family [7], were not detected. The OCG annotated with MYB12 contained 336 sequences, indicating that the boundaries within this gene family might be blurred, potentially leading to the merging of multiple MYB subgroup 7 members into a single OCG.

### Proof of concept: application to carotenoid biosynthesis

To further evaluate the performance of CoExpPhylo, we applied it to the carotenoid biosynthesis pathway, focusing on lutein and zeaxanthin branches. Using down-stream biosynthetic genes as input baits, the pipeline successfully detected multiple OCGs corresponding to known enzymes, as well as potential candidate genes involved in the pathway (Figure 4C).

Carotenoid biosynthesis starts from the precursor geranylgeranyl diphosphate (GGPP), which is converted to phytoene by phytoene synthase (PSY) [87]. Phytoene is subsequently desaturated by phytoene desaturase (PDS) and ζ-carotene desaturase (ZDS), forming lycopene [88, 89]. Lycopene cyclases, lycopene β-cyclase (β-LCY) and lycopene ε-cyclase (ε-LCY), catalyze the formation of α-carotene and β-carotene, key intermediates for xanthophyll biosynthesis [90]. Lutein biosynthesis proceeds via hydroxylation of α-carotene catalyzed by lutein deficient 5 (LUT5)/CYP97A3 and lutein deficient 1 (LUT1)/CYP97C1, while zeaxanthin is produced by hydroxylation of β-carotene, involving enzymes such as the β-carotene hydroxylases BCH1 and BCH2 [91–95].

#### Lutein

Focusing on the lutein branch of carotenoid biosynthesis, which plays a central role in photo-protection and light harvesting in plants, we evaluated CoExpPhylo’s performance using LUT1/CYP97C1 as bait. All known genes of the lutein pathway (PSY, PDS, ZDS, CRTISO, β-LCY, ε-LCY, LUT5/CYP97A3, and LUT1/CYP97C1) were recovered (Additional File 6), demonstrating the tool’s ability to reconstruct the main biosynthetic route.

No well-defined transcription factors could be identified in our analysis, as there is little consensus on transcriptional regulation of carotenoid biosynthesis across species and tissues [96]. Regulatory mechanisms are highly context-dependent, which makes detection through coexpression and phylogenetic approaches particularly challenging. Carotenoids, although sometimes associated with secondary traits such as pigmentation, are considered central metabolites due to their essential roles in photosynthesis and photoprotection, particularly in the case of lutein [97]. This underlines that while CoExpPhylo is particularly well-suited for specialized metabolism with more distinct pathway boundaries, it is also applicable to central metabolism, though in such cases, more stringent filtering strategies may be required.

#### Zeaxanthin

Focusing on the zeaxanthin branch, we recovered all known biosynthetic genes except CRTISO and BCH2, although both BCH1 and BCH2 were included as baits (Additional File 7). Interestingly, CoExpPhylo also identified LUT5/CYP97A3, LUT1/CYP97C1, and ε-LCY, which are primarily associated with lutein biosynthesis. This cross-detection likely reflects the fact that lutein biosynthetic enzymes are expressed constitutively across diverse conditions, causing their transcriptional profiles to co-vary with the more dynamically regulated zeaxanthin genes in our heterogeneous dataset.

Moreover, we detected a large number of clusters (706 OCGs), reflecting the fact that carotenoids, and especially xanthophylls, serve as precursors for a wide range of other compounds, including the plant hormones abscisic acid and strigolactones as well as various apocarotenoids [98, 99]. This metabolic branching makes distinguishing the core biosynthetic genes more challenging compared to specialized pathways.

### Tool benchmarking

CoExpPhylo allows the user to select either MAFFT [43, 44] or MUSCLE [45] for the multiple sequence alignment in Step 5. To evaluate the impact of the alignment method on the phylogenetic analysis, several PSS were processed using both tools. A representative dataset comprising 43 sequences revealed key differences between the alignments (Additional File 8). Five larger groups could be identified that contained the same sequences but were positioned differently and the internal branches within these groups were arranged differently. The preservation of major groupings shows that both tools are consistent with the overall evolutionary structure but differ in detailed phylogenetic distances, i.e. the connections between the groups are interpreted differently. These discrepancies stem from the different algorithms and scoring systems that both alignment tools use.

In contrast, a comparison of the different available tools for tree inference from these alignments – FastTree [46, 47], RAxML [48] or IQ-TREE [49] – revealed greater variability in sequence placement. Although the overall tree topology remained consistent, topological details varied across different tree-building approaches (Additional File 9). Notably, the phylogenetic differences introduced by alternative alignment methods exceeded those introduced by different tree inference tools.

The choice of tree inference software significantly influenced runtime performance (Figure 5). The utilization of MUSCLE instead of MAFFT for the global alignment step leads to a runtime increase of 9.5 minutes or 12.5 minutes, depending on the execution mode. Replacing FastTree with another phylogenetic tree inference tool has a stronger impact on the runtime performance: The utilization of IQ-TREE increased the computing time by over three and four hours, respectively. Executing MAFFT in combination with RAxML-NG takes over 140 hours, amounting to almost six days (Figure 5).

**Figure 5:**
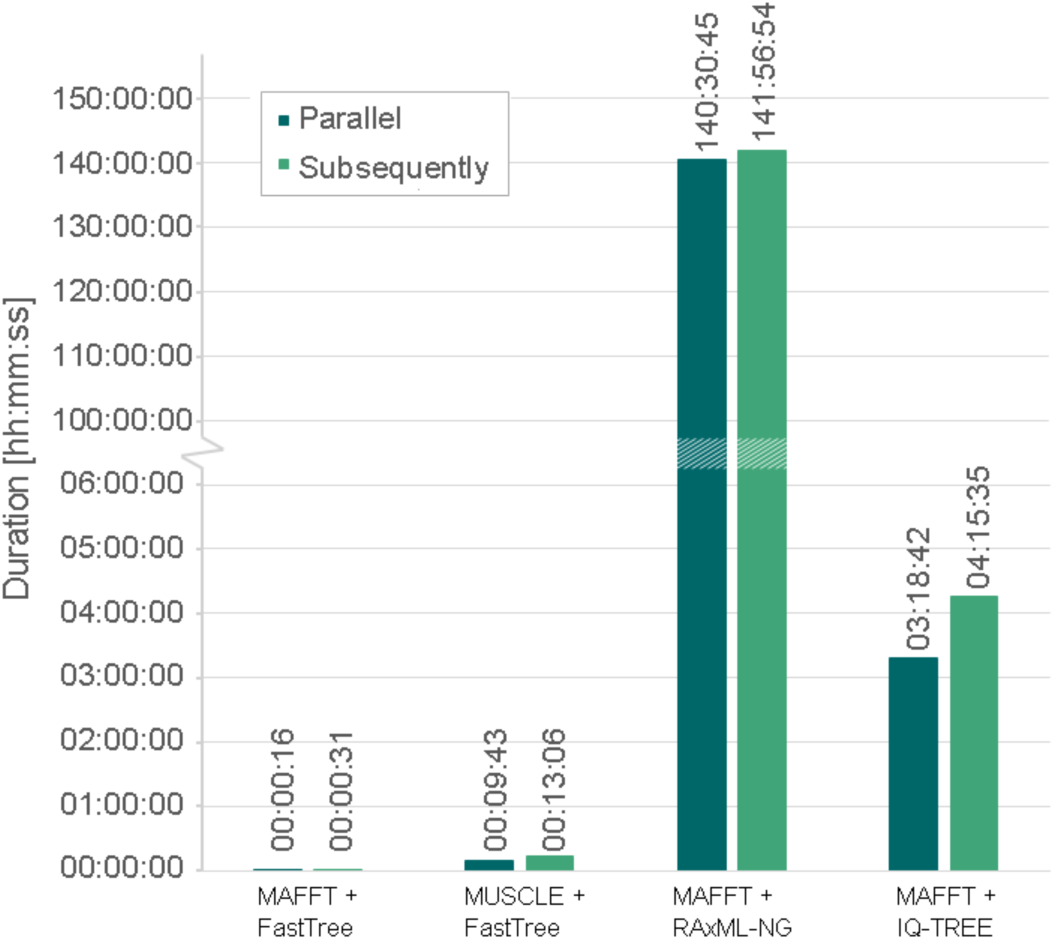
Comparison of the runtime performance of global alignment and phylogenetic tree construction. Shown are the runtime performances of the global alignment step combined with the construction of phylogenetic trees. Three representative sequence collections were selected (containing 43, 113, and 212 sequences) and processed in parallel (dark green) or subsequently (light green). The global alignment was conducted either with MAFFT v7.490 [43, 44] or MUSCLE v3.8.1551 [45], the phylogenetic tree was constructed with FastTree v2.1.11 [46, 47], RAxML-NG v1.2.2 [48], or IQ-TREE v2.0.7 [49]. The vertical axis was split between 6 and 100 hours in order to visualize all data points.

Parallel processing mitigated the impact of large OCGs on runtime. As all OCGs were processed simultaneously, the largest cluster determined the overall computation time. This effect was particularly pronounced for IQ-TREE and RAxML-NG, where the largest OCG (containing 212 sequences) accounted for 77.7% and 99% of the total runtime, respectively. Given that the dataset included 20 OCGs with over 1,000 sequences each, computational demands would increase substantially for larger datasets. Thus, the computational time would increase profoundly. Overall, a change in the global alignment method does not affect the runtime as severely as replacing FastTree by either IQ-TREE or RAxML-NG for these example OCGs. However, utilization of MAFFT on all OCGs of the used dataset does have a significant impact on the runtime as the global alignment of the three largest OCGs (> 2,000 sequences) took over 100 hours (data not shown).

In summary, the combination of MAFFT for global alignment and FastTree for phylogenetic tree construction represents the most efficient approach. Given that the biological results obtained with different methods show no substantial differences, but computational time increases dramatically with OCG size for alternative tools, the use of MAFFT and FastTree ensures an optimal balance between accuracy and performance.

## Conclusions

The discovery of multiple pathways based on a single bait gene in each indicates that CoExpPhylo can identify promising candidate genes of specialized biosynthetic pathways across multiple species, providing a solid basis for further functional and comparative analyses. By integrating coexpression analysis with phylogenetic relationships, the pipeline enables a broader, evolutionary-informed perspective on pathway organization and gene function. In addition to core enzymatic genes, CoExpPhylo can detect a range of transcription factors and transporters, offering a more comprehensive view of pathway regulation, even though it may not capture all regulatory players.

A key strength of CoExpPhylo is its ability to identify functionally related genes even in cases where coexpression pattern would be diffuse. This makes it a powerful tool for expanding pathway annotations beyond well-characterized model species. The generated OCGs offer a data-driven basis for highlighting candidate genes with potential involvement in a biosynthetic pathway, independent of prior pathway knowledge. However, these candidates should be considered as a starting point for further experimental validation. By offering a systematic and large-scale approach to gene discovery, CoExpPhylo helps to bridge the gap between computational predictions and functional characterization, addressing one of the major challenges in identifying novel genes with specific biochemical functions.

While CoExpPhylo effectively resolves many pathway components, opportunities for further refinement remain. For example, large gene families, such as MYB transcription factors, may exhibit high sequence similarity, leading to broader clustering in some cases. Furthermore, in some cases, genes from closely related metabolic pathways were grouped within the same OCGs. This highlights the need for additional functional validation to precisely determine pathway specificity, particularly for multifunctional enzymes or regulatory proteins. Additionally, annotation quality is dependent on the reference database used, which may limit the classification of genes absent in *Arabidopsis thaliana*. Expanding reference data to include multiple species and refining the clustering approach for highly homologous sequences could further enhance the resolution of functional gene groups. Furthermore, incorporating a protein-protein interaction predictor would enhance the functionality of the program for pathways that form a metabolon. Accurate protein-protein interaction predictions could underline the likelihood of physical interaction, thereby supporting the functional relevance of identified candidate genes. In such cases, integrating predicted protein-protein interactions would help to differentiate between genes that are merely co-expressed or homologous and those that may directly cooperate at the protein level within the same metabolic pathway.

Despite these considerations, CoExpPhylo represents a valuable tool for biosynthetic pathway exploration, enabling the identification of both conserved and lineage-specific genes. By facilitating candidate gene discovery across diverse plant species, it offers new opportunities to investigate specialized metabolism with a systematic and scalable approach.

## Availability and requirements

**Project name:** CoExpPhylo

**Project home page:** https://github.com/bpucker/CoExpPhylo

**Operating system(s):** Tested on Ubuntu but should work on all Unix-like operating systems

**Programming language:** Python, Bash

**Other requirements:** Python 3.0 or higher

**License:** GNU GPL

**Any restrictions to use by non-academics:** not applicable

## Abbreviations

3GT: Anthocyanidin-3-O-Glycosyltransferase
4CL: 4-Coumarate:CoA Ligase
ANR: Anthocyanidin Reductase
ANS: Anthocyanidin Synthase
arGST: Anthocyanin-related Glutathion S-Transferase
BAN: BANYLUS
BCH: Beta carotenoid hydroxylase
β-LCY: Lycopene β -cyclase
C4H: Cinnamate 4-Hydroxylase
CDS: Coding Sequence
CHI: Chalcone Isomersae
CHS: Chalcone Synthase
CRTISO: Carotenoid isomerase
CYP97: Cytochrome P450, family 97
DFR: Dihydroflavonol 4-Reductase
ε-LCY: Lycopene ε -cyclase
F3’5’H: Flavonoid 3’,5’-Hydroxylase
F3’H: Flavonoid 3’-Hydroxylase
F3H: Flavanone 3-Hydroxylase
FLS: Flavonol Synthase
GGPP: Geranylgeranyl diphosphate
LAR: Leucoanthocyanidin Reductase
LDOX: Leucoanthocyanidin Dioxygenase
LUT: Lutein deficient
NCBI: National Center for Biotechnology Information
OCG: Orthologous Coexpressed Group
OMT: O-Methyltransferase
PA: Proanthocyanidin
PAL: Phenylalanine Ammonia-Lyase
PDS: Phytoene desaturase
PSY: Phytoene synthase
SRA: Sequence Read Archive
TPM: Transcripts Per Million
TT: Transparent Testa
UFGT: UDP-Glucose:Flavonoid Glucosyltransferase
ZDS: ζ-carotene desaturase

## Declarations

### Ethics approval and consent to participate

Not applicable.

### Consent for publication

All authors have read and approved the final manuscript.

### Availability of data and materials

The authors declare that all datasets used in this paper are available online [51, 52] and the Python code is published on GitHub: https://github.com/bpucker/CoExpPhylo.

### Competing interests

The authors declare no competing interests.

### Funding

Not applicable.

### Authors’ contributions

NG and BP planned the study and wrote the software. NG conducted the bioinformatic analyses. NG and BP interpreted the results and wrote the manuscript.

## Acknowledgements

This work was supported by the BMBF-funded de.NBI Cloud within the German Network for Bioinformatics Infrastructure (de.NBI) (031A532B, 031A533A, 031A533B, 031A534A, 031A535A, 031A537A, 031A537B, 031A537C, 031A537D, 031A538A). We also thank all members of the research group Plant Biotechnology and Bioinformatics for discussion and support. Open Access funding enabled and organized by Project DEAL and the University of Bonn.

## Additional files

**Additional File 1**

Additional_File_1.csv

Title: Species used for analysis and their taxonomic ranking.

Description: Shown are all species and their respective order and family used in this analysis. CDS and TPM files were collected from two data publications [51, 52].

**Additional File 2**

Additional_File_2.pdf

Title: Distribution of OCG sizes at different similarity and length thresholds.

Description: Shown are the number of sequences per OCG and their occurrence at different similarity and length cutoffs. Starting from default values (A), the similarity cutoff was reduced to 75 (C) and 70 (E), whereas the length cutoff was reduced to 90 (B) and 80 (D).

**Additional File 3**

Additional_File_3.txt

Title: Functional annotation of OCGs – Anthocyanins.

Description: Shown are the results of CoExpPhylo with respect to anthocyanins. The analysis was based on two transcriptomic data sets [51, 52] with *DFR*, *ANS*, *Bz2*, and *TT19* as bait genes. Samples with a high expression (>5 TPM) of *LAR* and/or *ANR* were excluded. The analysis was conducted with default parameters. The functional annotations refer to Araport11 *A. thaliana* Col-0 reference polypeptide sequences and and corresponding information [56].

**Additional File 4**

Additional_File_4.txt

Title: Functional annotation of OCGs – Proanthocyanidins.

Description: Shown are the results of CoExpPhylo with respect to proanthocyanidins. The analysis was based on two transcriptomic data sets [51, 52] with *LAR* and *ANR* as bait genes. The analysis was conducted with default parameters. The functional annotations refer to Araport11 *A. thaliana* Col-0 reference polypeptide sequences and and corresponding information [56].

**Additional File 5**

Additional_File_5.txt

Title: Functional annotation of OCGs – Flavonols.

Description: Shown are the results of CoExpPhylo with respect to proanthocyanidins. The analysis was based on two transcriptomic data sets [51, 52] with *FLS* as the bait gene. The analysis was conducted with default parameters. The functional annotations refer to Araport11 *A. thaliana* Col-0 reference polypeptide sequences and and corresponding information [56].

**Additional File 6**

Additional_File_6.txt

Title: Functional annotation of OCGs – Lutein.

Description: Shown are the results of CoExpPhylo with respect to lutein. The analysis was based on two transcriptomic data sets [51, 52] with *LUT1/CYP97C1* as bait genes. The analysis was conducted with default parameters. The functional annotations refer to Araport11 *A. thaliana* Col-0 reference polypeptide sequences and and corresponding information [56].

**Additional File 7**

Additional_File_7.txt

Title: Functional annotation of OCGs – Zeaxanthin.

Description: Shown are the results of CoExpPhylo with respect to zeaxanthin. The analysis was based on two transcriptomic data sets [51, 52] with *BCH1* and *BCH2* as the bait genes. The analysis was conducted with default parameters. The functional annotations refer to Araport11 *A. thaliana* Col-0 reference polypeptide sequences and and corresponding information [56].

**Additional File 8**

Additional_File_8.pdf

Title: Phylogenetic trees retrieved through different alignment methods.

Description: Shown are phylogenetic trees of one selected OCG retrieved from CoExpPhylo annotated with Chalcone-isomerase like. The analysis was based on two transcriptomic data sets [51, 52] with *DFR*, *ANS*, *Bz2*, and *TT19* as bait genes. The global alignment was conducted with either MAFFT [43, 44] or MUSCLE [45]. The tree was inferred through FastTree [46, 47] and visualized in iTOL [50]. Different colors of strips next to the labels indicate major phylogenetic groups. Sequences marked by green colored labels were collected via coexpression analysis. Trees are midpoint rooted.

**Additional File 9**

Additional_File_9.pdf

Title: Phylogenetic trees retrieved through different alignment methods.

Description: Shown are phylogenetic trees of one selected OCG retrieved from CoExpPhylo annotated with Chalcone-isomerase like. The analysis was based on two transcriptomic data sets [51, 52] with *DFR*, *ANS*, *Bz2*, and *TT19* as bait genes. The global alignment was conducted with MAFFT [43, 44] The tree was inferred either through FastTree [46, 47], RAxML [48] or IQTREE [49] and visualized in iTOL [50]. Sequences marked by green colored labels were collected via coexpression analysis. Trees are midpoint rooted.

## Notes

https://github.com/bpucker/CoExpPhylo

https://doi.org/10.24355/dbbs.084-202409160820-0

https://doi.org/10.24355/dbbs.084-202501230512-0

### Competing Interest Statement

The authors have declared no competing interest.

### Summary of Updates

- Carotenoid biosynthesis added as proof of concept - Selection of cutoff values and parameter settings explained - Run time analysis included - Example dataset added on GitHub - Extended documentation

