## Supplementary figures and images for "CoExpPhylo – A Novel Pipeline for Biosynthesis Gene Discovery"

### Additional File 2

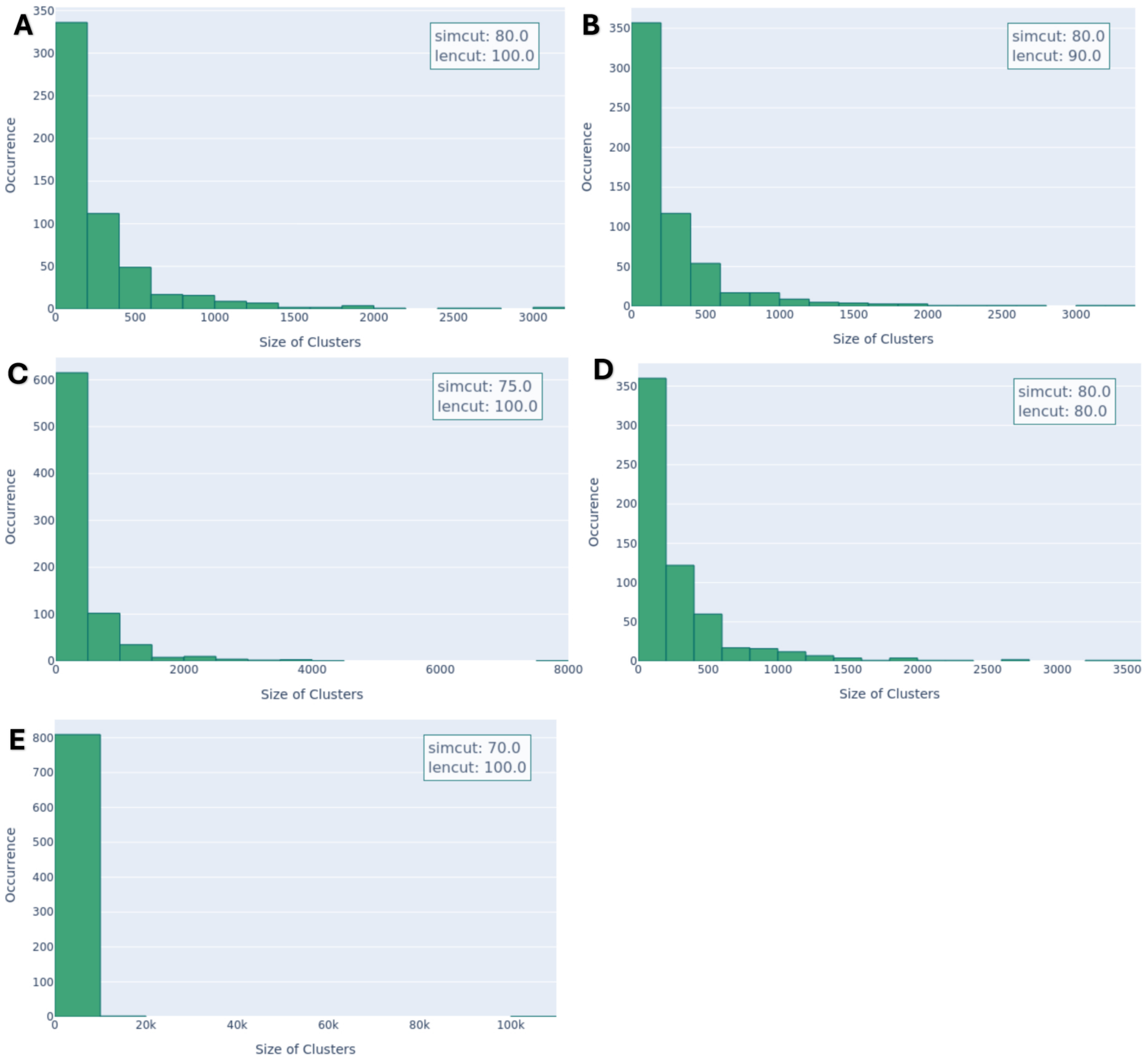

### Additional File 8

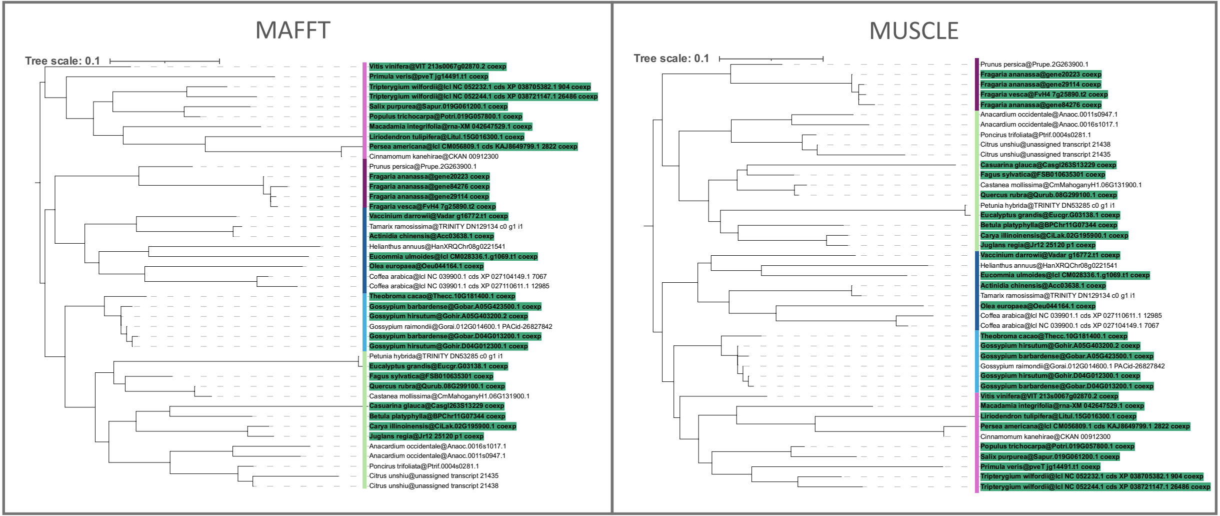

### Additional File 9

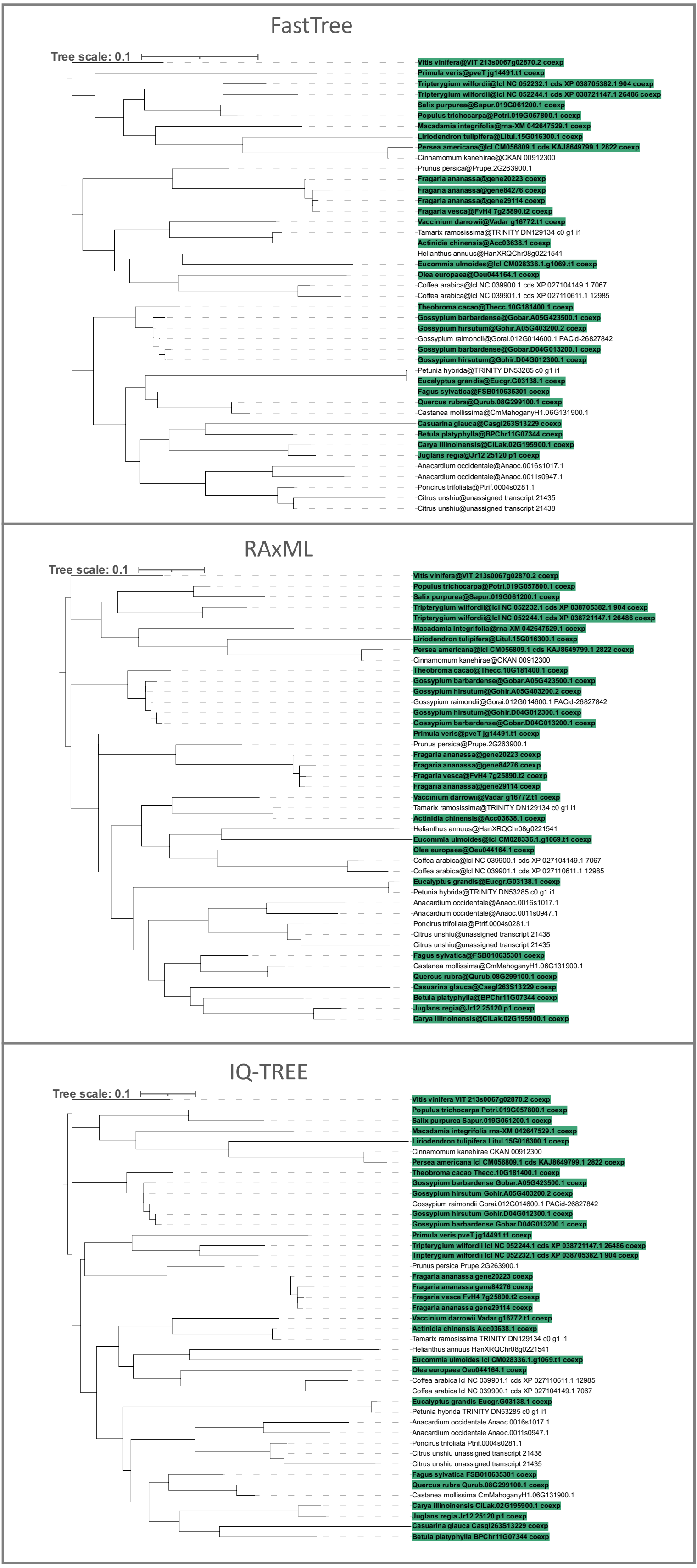
